# Hydraulic adjustments of Scots pine colonizing a harsh environment on volcano slopes

**DOI:** 10.1101/2023.02.22.528520

**Authors:** Têtè Sévérien Barigah, Fernanda Dos Santos Farnese, Paulo De Menezes Silva, Paul Humbert, Mustapha Ennajeh, Jérôme Ngao, Eric Badel, Hervé Cochard, Stephane Herbette

**Affiliations:** Université Clermont Auvergne, INRAE, PIAF, 63000 Clermont-Ferrand, France; Laboratory of Plant Stress Physiology, Instituto Federal de Educação, Ciência e Tecnologia Goiano, Campus Rio Verde, 75906□820 Rio Verde, GO, Brazil; Laboratory of Biodiversity and Valorization of Bioresources in Arid Zones, Faculty of Sciences of Gabes-City Erriadh, Zrig, Gabes 6072, Tunisia

**Keywords:** Acclimation, embolism, mortality, pioneer species, Scots pine, water stress

## Abstract

The ability of trees to survive and naturally regenerate in increasing drought conditions will depend on their capacity to vary key hydraulic and morphological traits that increase drought tolerance. Despite many studies investigating variability in these drought-tolerant traits, there has been limited investigation into this variability under recurrent severe drought conditions since the establishment phase.

We investigated the long-term hydraulic and leaf trait adjustments of Scots pine trees settled in an abandoned slag quarry by comparing them across three different topographic positions inducing contrasted effects on growth and development. We measured xylem and foliar traits to compare the water status of trees according to tree location and to evaluate the respective risk for xylem hydraulic failure using the soil-plant hydraulic model *SurEau*.

Compared to upslope and downslope trees, slope trees exhibited lower growth, vulnerability to embolism, specific hydraulic conductivity and photosynthetic pigment contents, as well as higher water potential at turgor loss point and midday water potentials. The hydraulic adjustments of trees settled on slag slopes reduced the risk for hydraulic failure and thus prevented an increase in embolism compared to downslope and upslope trees. These results suggest a prioritization of hydraulic safety over growth in Scots pine developed in a harsh environment, resulting in a dwarf phenotype.

## Introduction

Tree species with a wide distribution area or living in contrasting habitats, and pioneer species, must display variability for traits that allow them to cope with various environmental conditions. Especially, the soil water availability is critical in seedling establishment, plant growth and survival. Successful seedling and sapling development is a prerequisite for forest regeneration as well as for pioneer colonization (Lloret et al. 2009). Impeding these stages under climate change would hamper forest installation and regeneration (Anderson-Teixeira et al., 2013).

The phenotypic variability including plasticity and genetic variability is a key feature for tree species adaptation to environmental changes, especially for pioneer species (Sultan et al., 2000; Nicotra et al., 2010). Indeed, species with greater adaptive plasticity may be more likely to survive climate change that occurs too rapidly to allow for a population migration or an evolutionary response. Moreover, plasticity allows pioneer species to colonize environmentally diverse sites without the lag time required for local adaptation. Studying variations of traits related to drought resistance is necessary to understand the acclimation of pioneer species and to improve the predictions of tree responses to climate change (Anderegg, 2015).

Resistance to drought is conferred by a set of interacting traits, in which hydraulic traits have a key role (Martínez-Vilalta et al., 2002; Chaves et al., 2003; Choat et al., 2012). Indeed, drought-induced tree mortality is mainly due to xylem hydraulic failure caused by embolism (Barigah et al., 2013; Anderegg et al. 2016), even if functional evidence remains to be found to link hydraulic failure and tree death (Mantova et al., 2022). Much is known about the variability of these hydraulic traits across species (Maherali et al., 2004; Jacobsen et al., 2007; Choat et al., 2012), and about their intraspecific variability (e.g. in Martinez-Vilalta et al., 2009; Herbette et al., 2010; Bartlett et al., 2014; González-Muñoz et al., 2018). Within species variations in hydraulic traits have been reported for many species grown in natural conditions (Martínez-Vilalta et al., 2009; González-Muñoz et al., 2018), in common gardens (Wortemann et al., 2011; Lamy et al., 2011; Pritzkow et al., 2020) or under controlled experimental conditions (Awad et al., 2010; Lemaire et al., 2021). Yet, little is known about the physiological adjustment capacities of trees establishing under severe water deficit conditions over long periods. Studies comparing hydraulic properties along wide environmental gradients for a given species provide information on the extent of trait variations within the species’ range, but this did not reflect the effects of future more severe droughts. Conversely, most studies based on drought manipulation experiments submitted trees to severe drought but they were typically short term or did not consider the tree settlement phase (Cotrufo et al. 2011, MacKay et al. 2012, Herbette et al., 2021; Moreno et al., 2021). Yet, acclimation processes and feedback mechanisms could occur to adapt responses of trees to changes in water availability on longer time scales (Leuzinger et al. 2011, Barbeta et al. 2013, Feichtinger et al. 2014). For a full understanding of the drought resistance strategy of the tree species, the leaf traits need also to be considered, as they reflect the tree strategy regarding the water losses as well as the use of other resources (Wright et al. 2004, Bartlett et al. 2012). Moreover, trees can adjust their water consumption and hydraulic safety through the coordination of stomatal regulation and modification of the hydraulic system (Choat et al., 2012; Martínez-Vilalta et al., 2009; Rosas et al., 2019). Reflecting the stomatal control, the leaf water potential at turgor loss (*Ψ*_tlp_) and its related parameters have been proposed as a powerful indicator of the drought tolerance of plants (McDowell, 2011; Bartlett et al., 2012). Indeed, plants with lower *Ψ*_tlp_ are more efficient to maintain stomatal conductance, photosynthesis, hydraulic conductance, and growth under water stress. Despite stomatal closure, water loss continues through the cuticle and incompletely closed stomata, and this is measured as the leaf minimum conductance (*g*_min_). Recent studies point this trait as critical, especially for severe droughts (Duursma et al., 2019; Brodribb et al., 2020). In long-term drought conditions, a tight coordination between hydraulic and leaf traits should allow plants to balance between safety (avoiding hydraulic failure) and efficiency, in terms of resource use to ensure growth and maintenance (Brodribb et al. 2014). Therefore, to better understand pioneer settlement in harsh habitats and to predict forest development under a climate change, investigations are needed on the drought’s impact on hydraulic and leaf traits and over longer period starting from the seedling establishment.

Scots pine (*Pinus sylvestris* L.) spans a vast climatic gradient from Eastern Siberia to Southern Spain (Poyatos et al., 2007). It is a pioneer tree establishing on wet as well as on dry sites, on poor, sandy soils, rocky outcrops, peat bogs or other various harsh ecosystems (Richardson and Rundel, 1998). Although this species is considered to be drought resistant, more than one-third of the forest diebacks reported by Allen et al. (2010) are linked to Scots pine forests. These features make this species relevant for the study of the variability in hydraulic traits, in addition to economic reasons and its importance in successional dynamics of temperate forests and thus their sustainability. That’s why the variability of hydraulic traits has been studied in Scots pine (Poyatos et al., 2007; Martinez-Vilalta et al., 2009; Zang et al., 2012; Feichtinger et al., 2015; Seidel and Menzel, 2016, Rosas et al., 2019). However, more comprehensive surveys on hydraulic traits variability and their interactions are also needed in marginal populations to complete the picture of physiological and structural hydraulic plasticity in this pioneer species. This will contribute to improve the understanding of the colonization mechanisms of pioneer trees and help in predicting how climate change will affect this species.

We identified an abandoned quarry that has been exclusively colonized by Scots pines for 20 years, with no other vegetation that can settle, allowing to investigate adjustments mechanisms (throughout selection and acclimation) over many years under very constraining environmental conditions (Figure 1). The trees established on a slope displayed greatly reduced dimensions of plants, annual shoots and needles and yellowing of needles when compared to individuals established on downslope or upslope. In this study, we compared water relations of trees located in these different positions and we evaluated morphological and hydraulic traits at needle and stem levels that allowed this species to establish within such a harsh environment. We hypothesized that 1) Scots pine shows large difference between topographic conditions for both leaf and stem hydraulic traits including vulnerability to embolism, hydraulic conductivity, turgor loss point, stomatal conductance, leaf-to-sapwood area ratio, 2) the subsequent hydraulic adjustments in slope trees allow Scots pine trees to operate with a narrow hydraulic safety margin and at higher risk of mortality, 3) the slope trees display an embolism level below the lethal threshold of hydraulic failure and a minimal growth rate to renew structures.

**Figure 1:**
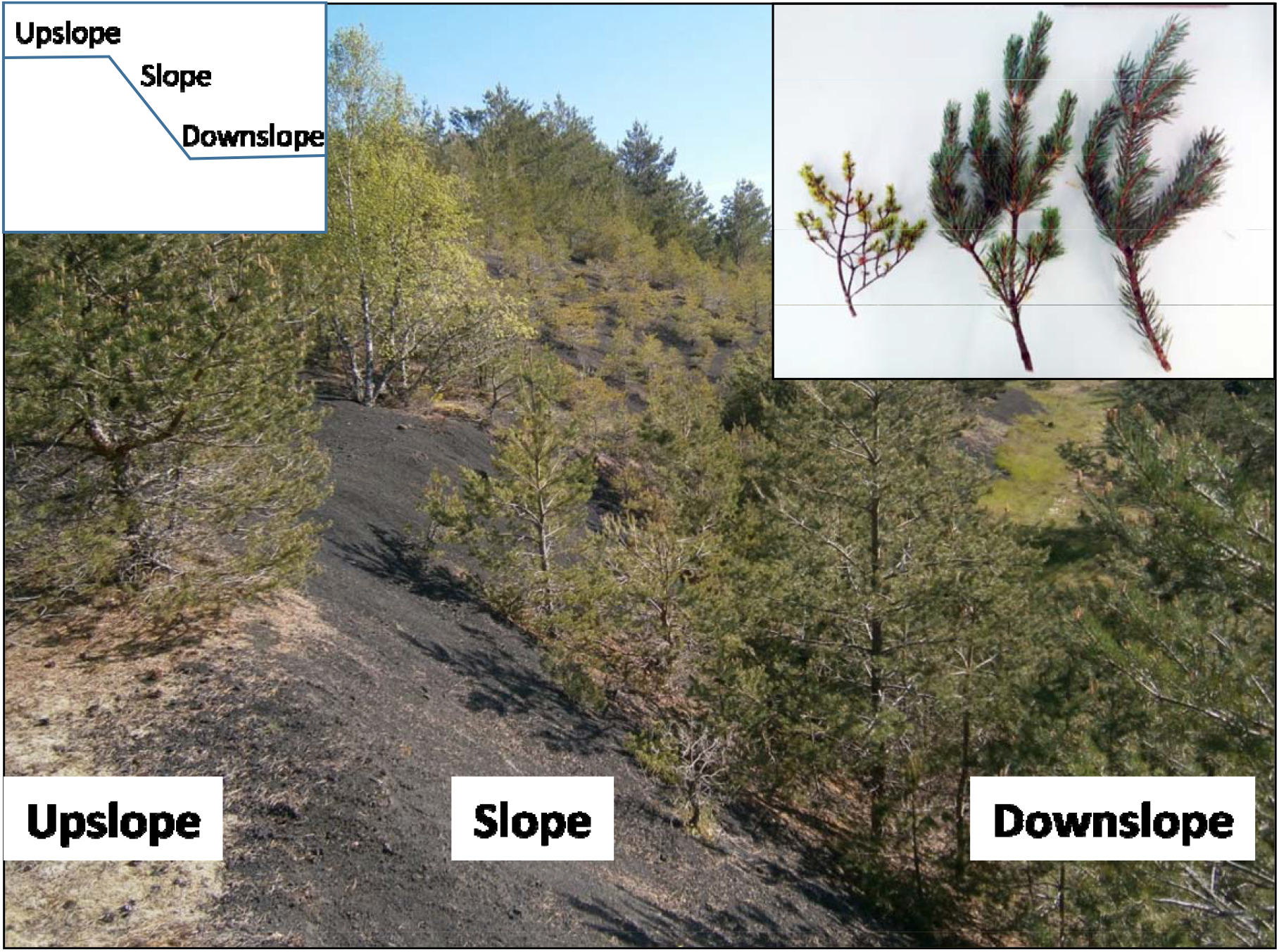
Study site and morphology of the studied trees. The study site is located on a former quarry of the volcano “Puy de la Vache”. The site is organized in terraces of slag separated by slopes. The vegetation present on these slopes and terraces has developed without any human intervention. On the slopes, the only vascular species to develop is the Scots pine. The site is split into 3 areas: the slope, the upslope (corresponding to a terrace) and the downslope (corresponding to the bottom of the quarry). The box shows the morphology of branches developed by slope, upslope and downslope trees, from left to right, respectively.

## Material and methods

### Site and trees

The site was located aside the East hillside of the volcano “Puy de la Vache”, (Saint-Genès-Champanelle, France), at an altitude of 962 to 973 m (45°42′ N, 2°58′ E), in a scoria quarry abandoned in the 1980’s. The soil substratum is scoria, a dark colored basaltic or andesitic volcanic rock. The site was characterized by loose substratum, low field capacity, nutrient poverty, high solar irradiation and temperature, recurrent summer droughts and cold winters (Frain, 1991). It was a former reserve of scoria and the company that operated the quarry shaped the landscape (Figure 1) leading to the removal of the soil, the slag substrate exposed and to shape the relief over a short distance in terraces, foothills and hillslopes.

On the site, Scots pines developed in a large open patch free of plant as the ground was left bare (Figure 1). We divided the studied trees in three groups regarding their location in the landscape and their statures. The slope trees were shrubby and small with reduced needle size, while the downslope trees were vigorous and tall with larger needles and the upslope ones were intermediate (figure 1). The selected trees were found to be homogeneous for age, from 17 to 23 years old, and an average of 18.8 years’ old whatever the group. Most of the measurements samplings were performed during the spring and summer of 2013. Some additional measurement such as the chlorophyll, carotenoids and mineral contents were performed during the summer 2018.

### Age and growth

The age of the trees, or the sampled branches, was assessed by the number of wood rings after coring the trunk or sampled branches.

Annual ring growth was measured on the branches collected for measurements of hydraulic traits and needle parameters. Mean annual ring growth was calculated as the total wood diameter over the number of rings for 31, 39 and 25 branches from downslope, upslope and slope trees, respectively.

### Needle chemical nutrient, chlorophyll and carotenoid content assays

We determined the contents of nitrogen (N), calcium (Ca), potassium (K), magnesium (Mg), sodium (Na), iron (Fe), copper (Cu), zinc (Zn) and plumb (Pb) on needles sampled from three to five trees per location. Needles were dried in a forced-air oven for 48 h at 80°C. Then, we determined the total N content using the Kjeldhal method, while we quantified Ca, K, Mg, Na, Fe, Cu, Zn and Pb by atomic spectrometric absorption method. First, we mineralized the samples of 20 mg of the dried needle material in 35% nitric acid and evaporated to dryness on a hotplate. We then dissolved the minerals in 0.1 N HCl solution. Afterward, we filtrated the solution on Whatman paper N°1. Finally, we measured the ion concentrations with an atomic absorption spectrometer (Avanta GBC spectrometer, Australia), using an air-acetylene flame. We reproduced this process thrice to evaluate a mean value for each sampled tree.

To assay the chlorophyll (Chl) and carotenoid contents of the needles, we relied on a procedure adapted from (Minocha et al., 2009) and sampled needles from 8 to 15 individual trees for each location and put them immediately in a cooler that contained crushed ice and brought them back to the laboratory. We transferred the samples in liquid nitrogen and stored them at -80 °C until analysis.

### Needle xylem water potentials

Six to eight trees per location were used for measurements of needle midday needle water potential (*Ψ*_md_) during the season of 2013, and five trees for daily time course of needle water potentials (*Ψ*_needle_) on August 21^st^, 2013. We completed the *Ψ*_md_ measurements between 11.00 am and 1.00 pm solar time. We processed on the field with a Scholander-type pressure chamber (PMS, Corvallis, Oregon, USA) on covered small shoot tips sampled from sunlight exposed south-facing shoots.

### Foliar traits

The specific needle area (SLA) was calculated as the needle area divided by its dry weight. The needle area was evaluated using the length and diameters and assuming the needle shape as a cone. We measured the SLA on 10 needles per tree and on five trees from each location.

The Huber value (*H*_v_) is the ratio of cross-sectional sapwood area to subtended leaf area, and it can therefore be analyzed as the ratio of hydraulic and mechanical investment costs over the expected gains obtained by leaf display. Needles from terminal branches were oven-dried and weighed, and SLA was used to convert the total dry weight of the distal needles of each branch into total branch needle area. Sapwood area was estimated by measuring the total xylem area on digital images of scanned cross-sections using ImageJ software (Wayne Rasband-National Institute of Health).

The needle-cell turgor features were computed using pressure-volume curves (p-v curves). The bench dehydration method was used (Tyree and Hammel 1972; Sack and Pasquet-Kok, 2010). We cut 50 cm long shoot samples from eleven to thirteen trees per location and immediately placed them in plastic bags with wet paper towels to bring them in the laboratory. The samples were rehydrated at full turgor by putting the cut end in water at room temperature overnight in a sealed bag and in a dark room. Then, shoots bearing needles were allowed to dehydrate while periodically being weighed and measured for water potential (*Ψ*) using a pressure chamber (1505D, PMS Instrument). To prevent rapid water loss, the shoot samples could be kept in plastic bags between measurements. When *Ψ* was lower than - 4 MPa, we measured the needle area, and the entire shoot were dehydrated in an oven at 70 °C during 72 h to reach the needle dry weight. The p-v curve was plotted using the inverse values of needle water potential versus the relative water content (RWC) which was obtained from repeated determinations of fresh mass during dehydration of the shoot tips. Several parameters can be extracted from the p–v curves (Tyree and Hammel, 1972, Sack and Pasquet-Kok, 2010) : the osmotic potential at full turgor (π_0_) and the water potential at turgor loss point (*Ψ*_TLP_), the water content at full turgor (*WC*_0_) calculated as the mass of water per needle dry mass, the relative water content at turgor loss point (*RWC*_TLP_) and the bulk modulus of tissue elasticity (ε_0_).

The minimal needle conductance (*g*_min_) was also determined using data from p-v curves (detailed above). The *g*_min_ was determined from the slope of the linear part of the curve after stomatal closure, and adjusted for needle area of the sample and for temperature and humidity regularly scored during measurements.

### Hydraulic traits

To visualize xylem embolism, we relied on X-ray microtomograph observations, which is a reference method (Cochard et al., 2015). We excised 0.03m-long xylem segments from branches of two trees per location. We removed the needles under water and sealed them in liquid paraffin wax in order to prevent dehydration during the X-ray scans. We placed the samples in an X-ray microtomograph (Nanotom 180 XS, GE, Wunstorf, Germany). The field of view was 5 × 5 × 5 mm and covered the full cross section of the samples. The X-ray source settings were 50 kV and 275 μA. For every 40-min scan, we recorded 1000 images during the 360° rotation of the sample. After 3D reconstruction, the spatial resolution of the image was 2.5 × 2.5 × 2.5 μm per voxel. We extracted the transverse 2D slice from the middle of the volume and used ImageJ software for visualization and image analysis.

Vulnerability to embolism was measured on branches from at least five individuals per location. Branches of 0.4–0.5 m long were harvested, wrapped in humid paper and plastic bag, brought to the laboratory and stored at 4°C until measurements in the following days. We cut stem segments of 0.28 m-long just before the measurement of their vulnerability to cavitation with the Cavitron technique (Cochard et al., 2005). We measured the loss of hydraulic conductance of the stem segment while the centrifugal force generated an increasing negative xylem pressure. The plot of the percent loss of xylem conductance (PLC) versus the xylem pressure represents the vulnerability curve. First, we set the xylem pressure to a reference pressure (−0.5 or –1.0 MPa) and we determined the maximal conductance (*k*_max_) of the sample. Then, we set the xylem pressure to a more negative pressure for 2 min, and we determined the new conductance (*k*). The procedure was repeated for more negative pressures (with –0.50 MPa increments) until PLC reached at least 90%. We generated vulnerability curves for each branch by fitting the data with exponential–sigmoidal function (Pammenter and Willigen, 1998):

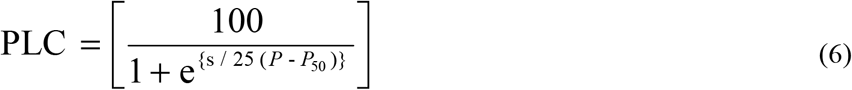

Where *P*_50_ is the pressure causing 50% loss of conductance and s is the slope value at this point.

Specific hydraulic conductivity (*K*_s_) was measured on branches from six individual trees per location. Branch segments were recut underwater using a razor blade and their length (*L*_stem_) was measured. The apical end of the sample was plugged to an embolism meter (Xyl’em, Bronkhorst, Montigny les Cormeilles, France). The total conductance (*K*) was then measured under low pressure (2–7 kPa) using a solution of 10 mM KCl and 1 mM CaCl_2_. The xylem area *A*_x_ was measured as the mean area of the both ends of the sample on digital images of scanned cross-sections in ImageJ. The *K*_s_ was defined as follows:

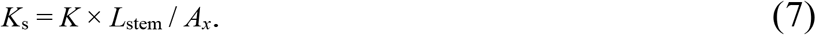

Leaf specific conductivity (LSC) was measured on the same samples and was determined by replacing Ax to the subtended leaf area. This leaf area was measured as previously described.

### SurEau model simulations

The SurEau model is a mechanistic discrete-time soil–plant–atmosphere hydraulic model that predicts plant hydraulic and hydric properties under simulations of water stress (Martin St Paul et al., 2017; Cochard et al., 2021). It is used to simulate the time to hydraulic failure under drought considering that this time is reached with high embolism level, i.e. 90 % (e.g. Lemaire et al., 2021). In this work, we used the SurEau model to understand how the combination of tree traits measured in the 3 quarry ecological conditions impacted their time to reach hydraulic failure. We parameterized the trees of each condition in the model by taking into account their average size, estimated in the field, and the physiological parameters that we measured (see the parameter table in the Supplementary data at *Tree Physiology* Online for an exhaustive list of variables). For the soil, we considered that the surface explored by the roots was the same as the surface of the crowns and that the ratio between the volume of water available in the soil and the leaf surface was constant. This defined the soil depth in each condition. The hydric and hydraulic properties of the soil were described by pedotransfer functions according to the model of van Genuchten (1980) adapted to scoria (Wallach et al 1992). We considered stomatal conductance to be a linear function of needle turgor pressure calculated from the pressure-volume curve parameters. The water potential at stomatal closure was assumed to coincide with the turgor loss point. The bark conductance was assumed to be equal to the minimum conductance measured on leaves. The exhaustive list of variables can be found in the table S1 at Tree Physiology Online. The model was run under constant climate conditions set to match the climate conditions in the field at noon on a typical sunny day in June, with the atmospheric temperature set at 28 °C, the atmospheric relative humidity set at 25% and the PAR set at 1000 μmol m−2 s−1. The wind speed was set at 1 m s−1. In a first case, simulations are made for each type of tree in their respective environments. The combined effects of morphological and physiological parameters on hydraulic failure are then taken into account. In the next two cases, we simulated trees in a downslope and slope position but applied the physiological traits measured in the other site conditions. Here, the morphological differences between the conditions are ignored. The results of the simulations are given in terms of the temporal dynamics of PLC of the branches.

### Statistical analysis

We subjected measured and derived data to statistical analysis using a software package (XLSTAT 19.4.46926, Addinsoft, Paris, France) for ANOVA. We compared the mean values with Tukey’s multiple range tests at 0.05 levels when effects were significant.

## Results

For investigating foliar and hydraulic traits, we sampled branch samples of homogeneous dimensions. The samples showed differences in age, especially between downslope and upslope trees (figure 2). The slope trees showed a very weak annual ring growth rate over years (P<0.001) compared to downslope and the upslope trees. To test if slope, even upslope, trees suffered from nutrient deficiency, we measured the main chemical elements of the needles (table 1). Except for nitrogen, there was no difference in needle nutrient contents between the locations. Both upslope and slope trees displayed similar nitrogen content (8.54 ± 0.35 and 7.16 ± 0.49 mg g^-1^ DW), contrary to the downslope trees that had higher content of nitrogen (12.85 ± 1.03 mg g^-1^ DW). The total needle chlorophyll and carotenoid contents were also variable regarding the tree locations (table 2), They were significantly higher in the downslope trees compared to the slope and upslope trees. To compare the water status of trees according to their position, we monitored the evolution of needle midday water potential (*Ψ*_md_) over the growing season (figure 3A). At the onset of the experiment, we found no significant difference in *Ψ*_md_ between locations. Then, the *Ψ*_md_ decreased over the season but stabilized earlier for the slope trees than for the upslope and downslope ones. Consequently, the slope trees had a significantly higher *Ψ*_md_ (about – 1.0 MPa) compared to the upslope and downslope ones (up to -1.5 MPa). The *Ψ*_md_ were similar for both upslope and downslope trees (figure 3A). On day 233, sunny and driest day in late summer of the year, we carried out a survey of needle water potentials (*Ψ*, figure 3B). At predawn (i.e. before 7 am), no significant difference in *Ψ* was found between trees from the three locations. Then, the *Ψ* decreased over the time but stabilized earlier for the slope trees than for the upslope and downslope ones. Consequently, the slope trees had a significantly higher *Ψ* compared to the upslope and downslope ones, during this day (figure 3B). We found similar results for day 214, which was another almost cloudless and driest day in the same summertime (data not shown).

**Table 1:** Nutrient content in needle from Scots pines developed under the three conditions. Data are means values (± SE) from three branches per tree sampled from three to five trees developed on the downslope, the upslope and the slope, respectively. Different letters indicate significant differences between the conditions for each element (*P* < 0.05).

| Element Content (mg.g<sup>-1</sup> DW)) | Downslope | Upslope | Slope |
| --- | --- | --- | --- |
| Azote | 12.85(± 1.03)<sup>a</sup> | 8.54 (± 0.35)<sup>b</sup> | 7.16 (± 0.49)<sup>b</sup> |
| Potassium | 7.64 (± 0.55) | 6.89 (± 0.29) | 7.50 (± 0.38) |
| Calcium | 14.3 (± 0.79) | 13.4 (± 0.76) | 14.7 (± 0.39) |
| Magnesium | 4.65 (± 0.22) | 4.87 (± 0.12) | 5.49 (± 0.33) |
| Iron | 0.451 (± 0.027) | 0.514 (± 0.062) | 0.494 (± 0.042) |
| Copper | 0.0492 (± 0.0026) | 0.0415 (± 0.0054) | 0.0416 (± 0.0033) |
| Zinc | 0.0552 (± 0.0021) | 0.0450 (± 0.0058) | 0.0407 (± 0.0030) |

**Table 2:** Photosynthetic pigment content in needle from Scots pines developed under the three conditions. Data are means values (± SE) from 11, 8 and 15 individual trees developed on the downslope, the upslope and the slope, respectively. Different letters indicate significant differences between the conditions for each pigment content (*P* < 0.01).

| Pigment Content | Downslope | Upslope | Slope |
| --- | --- | --- | --- |
| Total chlorophyll (mg.g<sup>-1</sup> FW) | 1.98 (± 0.20)<sup>a</sup> | 0.99 (± 0.08)<sup>b</sup> | 0.58 (± 0.04)<sup>c</sup> |
| Chlorophyll a/b ratio | 2.18 (± 0.09)<sup>a</sup> | 2.28(± 0.06)<sup>a</sup> | 2.55(± 0.03)<sup>b</sup> |
| Total carotenoids (mg.g<sup>-1</sup> FW) | 0.511 (± 0.049)<sup>a</sup> | 0.261 (± 0.021)<sup>b</sup> | 0.190 (± 0.010)<sup>c</sup> |

**Figure 2:**
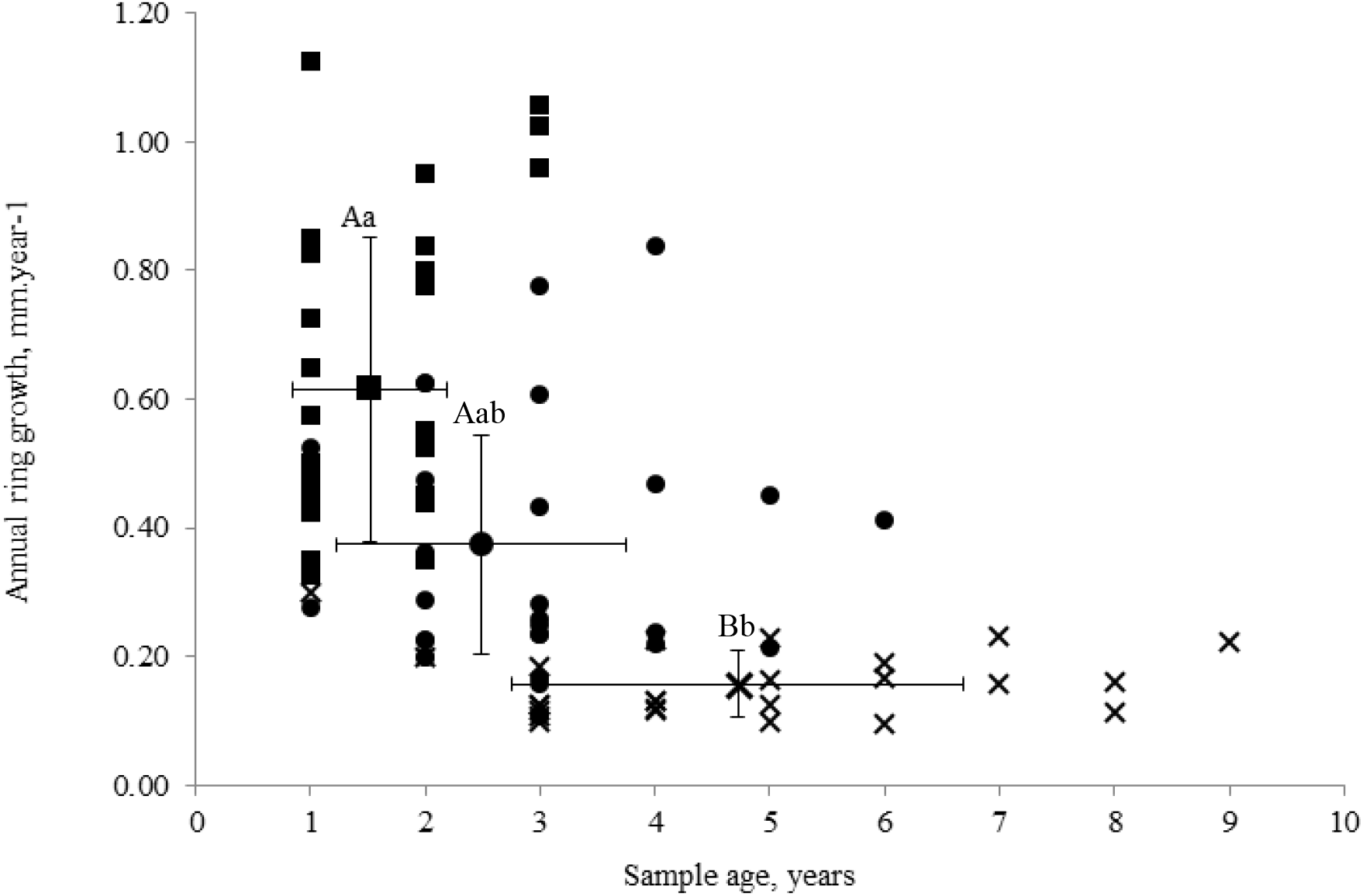
Annual ring growth of branch samples from Scots pines developed under the three conditions. Branches were sampled on trees located on the downslope (square), the upslope (circle) or the slope (cross). Their annual ring growth were measured and plotted versus their age. Each point represents one branch from one individual tree. Points showing error bars (± SD) are mean values, and different uppercase and lowercase letters indicate significant differences between conditions for annual ring growth and sample age, respectively (P < 0.05).

**Figure 3:**
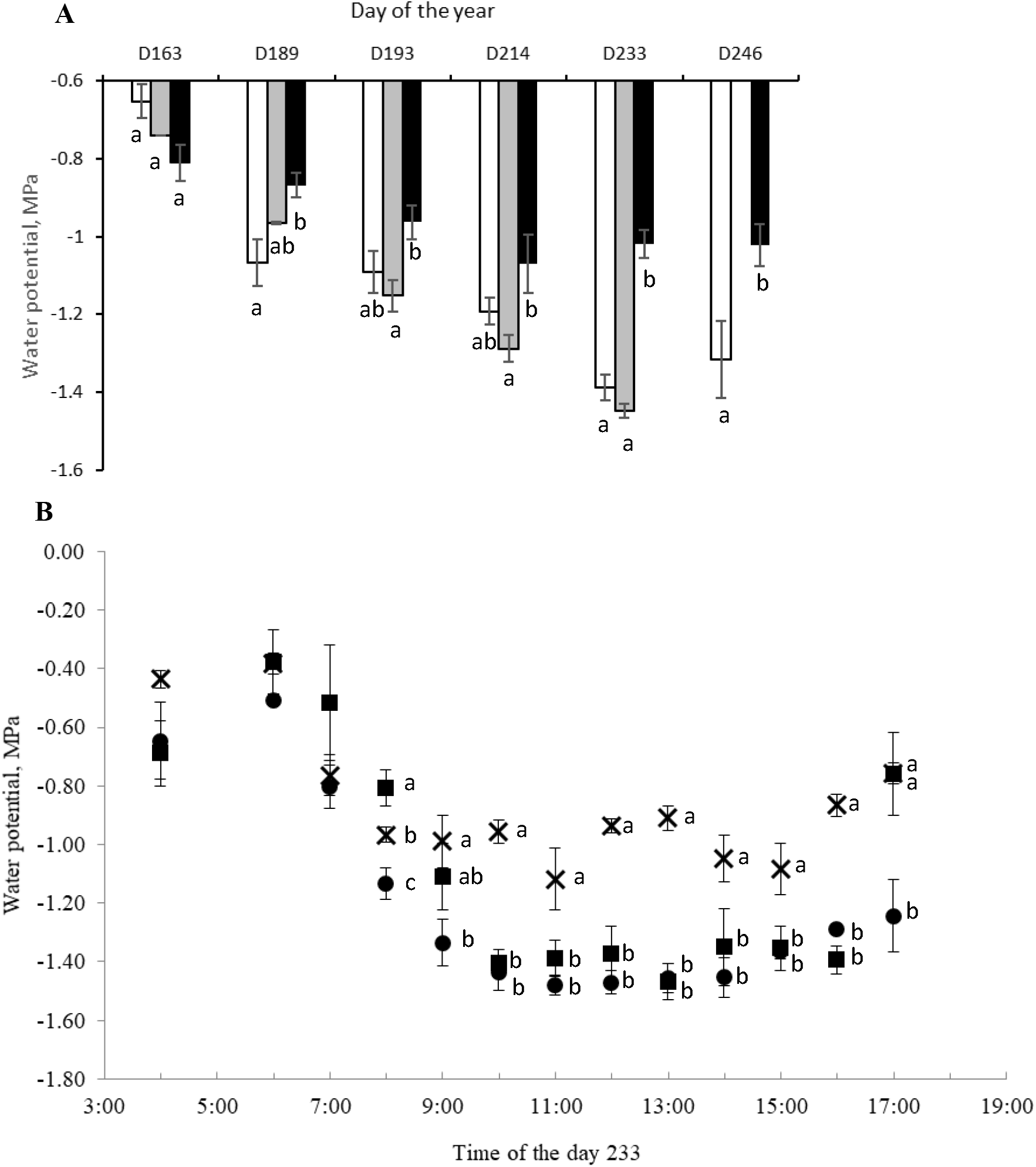
Seasonal variations (A) and daily time course (B) in xylem water potential of Scots pines developed under the three conditions. A, Midday water potentials were measured on tips of branches sampled from five trees located on the downslope (white), the upslope (grey) or the slope (dark), during sunny days preceded by a period of several days without rain. B, Water potentials were measured regularly from predawn (4 am) to the end (5 pm) of the day 233 on branches sampled from 5 trees located on the downslope (square), the upslope (circle) or the slope (cross). Symbols are mean values, bars represent standard errors, and different letters indicate significant differences between the conditions for each time (*P* < 0.05).

To investigate the drought resistance strategy according to the tree’s location, we measured hydraulic and foliar traits. We measured the xylem vulnerability to embolism on branches (figure 4 A). The slope trees were the most resistant to embolism with a branch *P*_50_ value of -3.70 MPa, while the downslope trees were the most sensitive with a branch *P*_50_ value of -3.25 MPa (figure 4A, table 3). The X-ray microtomography observation revealed that the native embolism in the tested trees was rather low (less than 5 %) regardless their location, and no embolism did occur over the last years (figure 5). This excludes possible effect of native embolism on the measured *P*_50_. The branches from slope trees showed also a significantly lower mean value for *K*_s_ and *LSC* compared to upslope and downslope trees (table 3).

**Table 3:** Functional traits for Scots pines developed under the three conditions. Data are the mean values (± SE) and different letters indicate significant difference between the conditions for each trait (*P* < 0.05). ε_0_, bulk modulus of tissue elasticity; g_min_, minimal needle conductance; *H*_v_, Huber value ; *K*s, specific hydraulic conductivity of the branch ; *LSC*, leaf specific conductivity of the branch; *P*_50_, pressure inducing 50 % loss of hydraulic conductance; Ψ_*TLP*_, water potential at turgor loss point; π_0_, osmotic potential at full turgor; RWC_*TLP*_, relative water content at turgor loss point *SLA*, specific leaf area ; *WC*_0_, water content at full turgor.

| Traits (units) | Downslope | Upslope | Slope |
| --- | --- | --- | --- |
| $P_{50 \text{ stem}}$ (MPa) | -3.25 ($\pm 0.05$)<sup>a</sup> | -3.49 ($\pm 0.11$)<sup>ab</sup> | -3.70 ($\pm 0.07$)<sup>b</sup> |
| $K_s$ (mol m<sup>-1</sup> s<sup>-1</sup> MPa<sup>-1</sup>) | 23.3 ($\pm 2.41$)<sup>a</sup> | 24.4 ($\pm 2.11$)<sup>a</sup> | 11.5 ($\pm 1.06$)<sup>b</sup> |
| $LSC$ (mmol m<sup>-1</sup> s<sup>-1</sup> MPa<sup>-1</sup>) | 5.29 ($\pm 0.38$)<sup>a</sup> | 5.96 ($\pm 0.49$)<sup>a</sup> | 3.14 ($\pm 0.41$)<sup>b</sup> |
| $SLA$ (m<sup>2</sup> g<sup>-1</sup>) | 80.79 ($\pm 1.14$)<sup>a</sup> | 67.75 ($\pm 1.48$)<sup>b</sup> | 108.94 ($\pm 10.24$)<sup>c</sup> |
| $H_v$ (cm<sup>2</sup>.m<sup>-2</sup>) | 2.62 ($\pm 0.18$) | 2.70 ($\pm 0.14$) | 2.64 ($\pm 0.11$) |
| $g_{\min}$ (mol m<sup>-2</sup> s<sup>-1</sup>) | 1.91 ($\pm 0.23$)<sup>a</sup> | 2.79 ($\pm 0.17$)<sup>b</sup> | 1.64 ($\pm 0.19$)<sup>a</sup> |
| $\Psi_{TLP}$ (MPa) | -2.07 ($\pm 0.11$)<sup>a</sup> | -1.87 ($\pm 0.15$)<sup>ab</sup> | -1.78 ($\pm 0.10$)<sup>b</sup> |
| $WC_0$ | 2.04 ($\pm 0.44$) | 1.67 ($\pm 0.30$) | 1.57 ($\pm 0.24$) |
| $RWC_{TLP}$ | 70.76 ($\pm 5.91$) | 75.21 ($\pm 4.18$) | 76.14 ($\pm 3.84$) |
| $\pi_0$ | -1.36 ($\pm 0.10$) | -1.32 ($\pm 0.11$) | -1.21 ($\pm 0.08$) |
| $\varepsilon_0$ (MPa) | 6.28 ($\pm 1.65$) | 5.68 ($\pm 1.12$) | 5.11 ($\pm 1.01$) |

**Figure 4:**
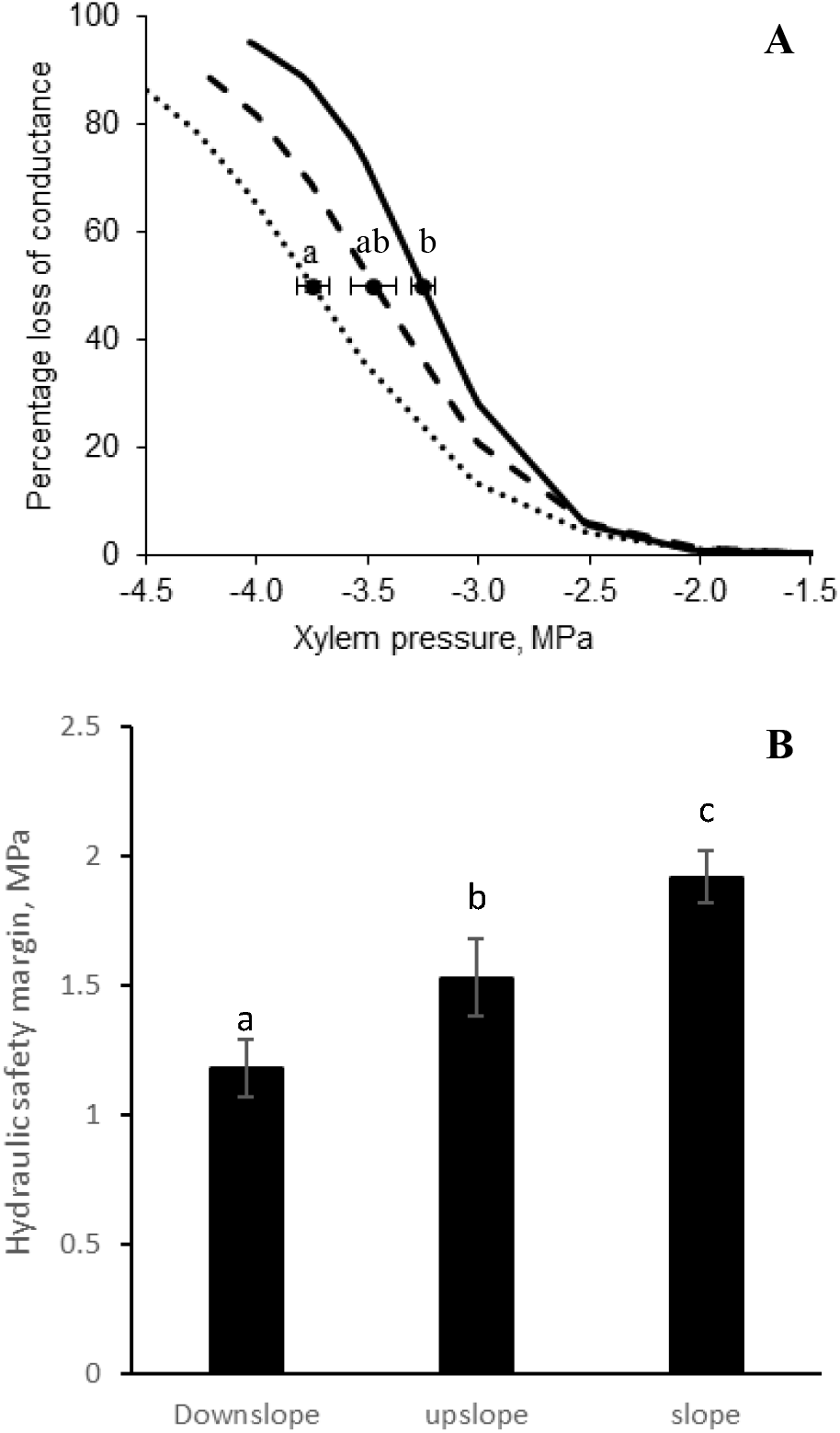
Xylem vulnerability to embolism (A) and hydraulic safety margin (B) of branches from Scots pines developed under the three conditions. A, vulnerability curves were performed from measurements on branches sampled from 6 trees located on the downslope (solid line), on the upslope (dashed line) or on the slope (dotted line). For each curve, the mean water potential at 50% loss of hydraulic conductance (*P*_50_) is indicated by the open symbol and the error bar is the standard error. B, the hydraulic safety margin was calculated for each tree from the difference between and *Ψ*_TLP_ and *P*_50_ values. Mean values and standard errors are represented. Different letters indicate significant differences between the conditions (*P* < 0.01).

**Figure 5:**
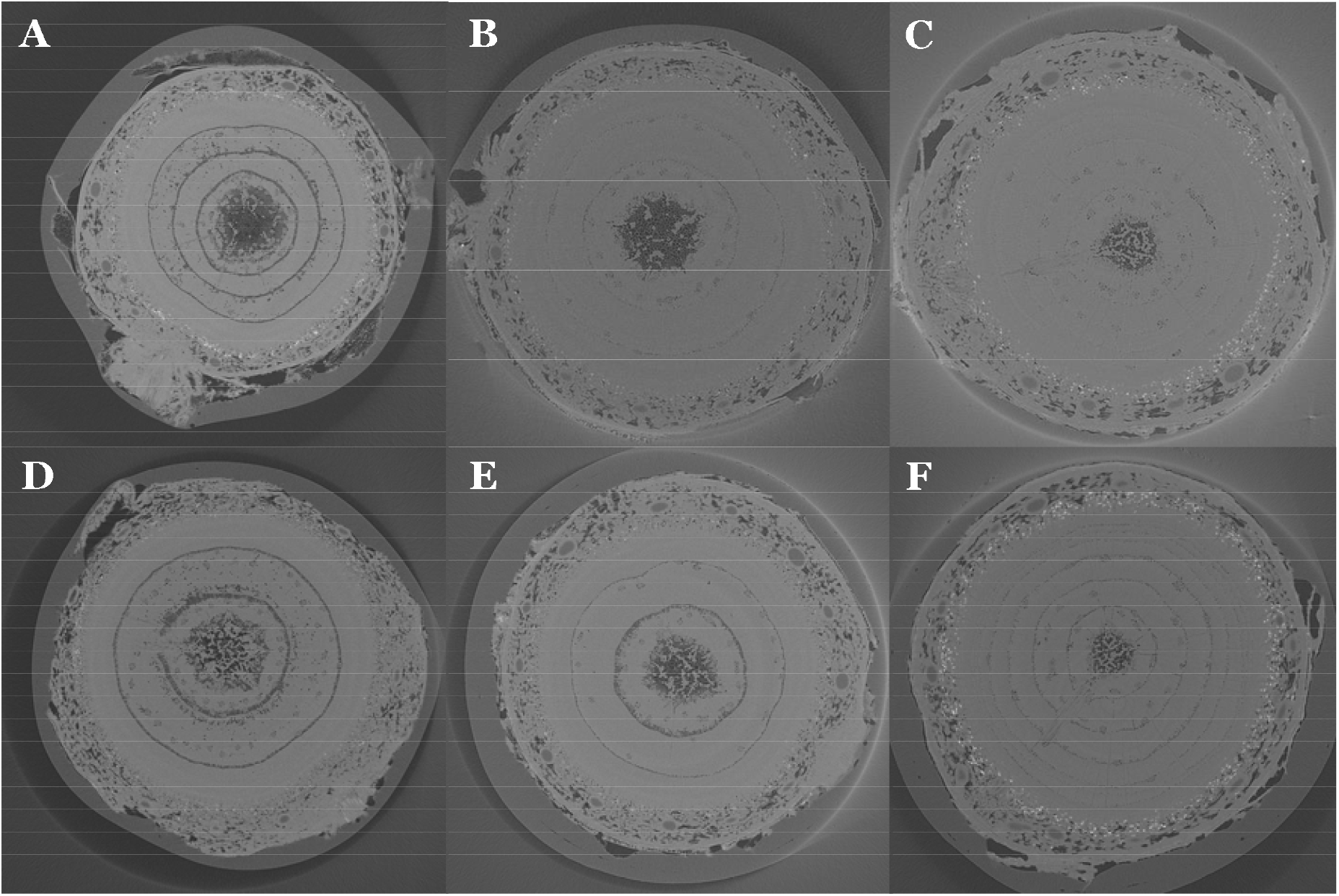
Transverse X-ray microtomography images of branches from Scots pines developed under the three conditions. Branches were sampled from trees developed at the downslope (A, D), at the upslope (B, E) or at the slope (C, F). Embolized tracheids are as depicted by black color while the fully saturated tissues are depicted as bright color. Embolized conduits were mainly found for the oldest wood rings. No embolism (or few) was observed in the rings of the last two years.

At the needle level, the SLA showed significant differences according to the tree location, the SLA being significantly higher in slope trees compared to downslope and upslope trees. The bulk leaf parameters drawn from the p-v curves revealed significant differences *Ψ*_TLP_ between tree locations (table 3), whereas other related parameters including WC_0_, RWC_TLP_, π_0_ and ε_0_ showed some differences but not significant, especially between the downslope and the slope trees. The *g*_min_ were also different between the tree locations, the upslope trees showing the greatest mean values (table3). Although, there were large differences in needle dimensions (figure 1) according to the tree locations, the *H*_v_ were similar between tree groups (table 3).

The risk for xylem hydraulic failure according to the location was evaluated in two ways: i) by calculating the hydraulic safety margin, ii) by simulating with the SurEau model the impact of a drought on the dynamics of xylem embolism. We calculated the hydraulic safety margin as the difference between *P*_50_ and the *Ψ*_TLP_. The former is considered as a threshold in hydraulic failure for conifers, while the latter is a proxy of the water potential inducing stomatal closure delaying embolism occurrence. The figure 4B shows that this safety margin was significantly higher for slope trees (mean of 1.92 MPa) compared to downslope trees (1.18 MPa), upslope trees having an intermediate mean value (1.53 MPa). Using the experimental trait values and the pedo-climatic conditions, the computations of the SurEau model enlightened that the PLC dynamics showed differences between tree locations (figure 6). In their respective locations, xylem embolism increased later in downslope trees comparing to slope and upslope trees (figure 6A). finally, there was no difference in xylem embolism dynamic between slope and upslope trees. However, when trees were submitted to similar drought conditions, i.e. drought on downslope or on slope (Figure 6B and C, respectively), hydraulic failure was delayed in trees grown on slope compared to trees grown on upslope or on downslope.

**Figure 6:**
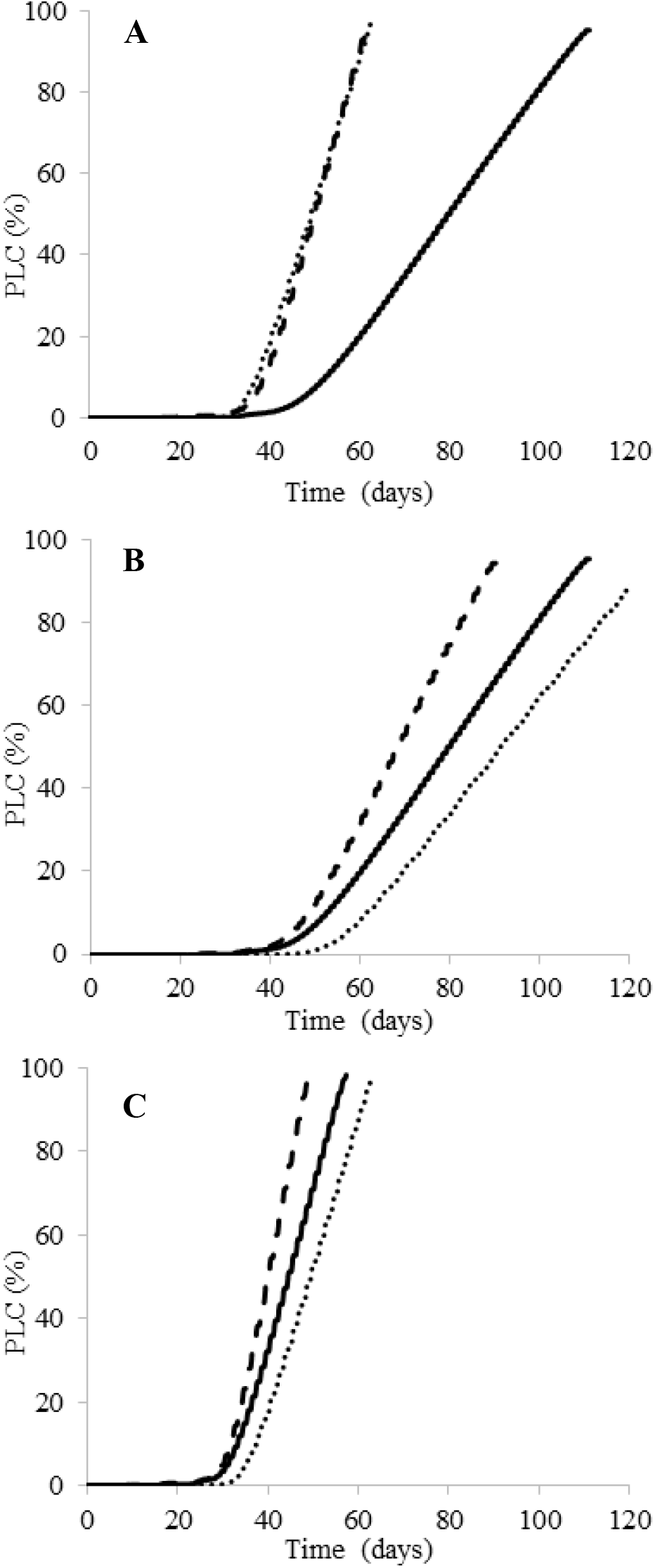
Simulated (SurEau model) time courses of the loss of xylem hydraulic conductance in the branches during a virtual drought event. Drought simulations were performed using experimental traits values from trees developing on the downslope (solid lines), on the upslope (dashed lines) or on the slope (dotted lines), while considering three conditions: trees in their respective locations (A), all trees on the downslope (B) and all trees on the slope (C).

## Discussion

Scots pines settled and developed under the most severe stress conditions, i.e. on slopes, were those with the greatest hydraulic safety margin and were operating with the highest water potentials, to avoid embolism events. This hydraulic safety seems to be prioritized over growth, which is strongly reduced. These trees also presented a reduced photosynthetic capacity, with a reduced hydraulic conductivity, a low quantity of photosynthetic pigments, and bulk leaf parameters such as π_0_, *Ψ*_TLP_, and ε_0_ unfavorable to gas exchanges. Scots pines in intermediate stress conditions, i.e on upper slope, showed intermediate hydraulic adjustments.

The physiological and morphological adjustments in Scots pines settled and developed over a long time (about 20 years) and under very stressful conditions were quite different from those observed for the same species between populations located along environmental gradients throughout Europe (Martinez-Vilalta et al., 2009; Rosas et al. 2019). For instance, we observed a decrease in stem conductivity and vulnerability to embolism, no change in *H*_v_ and an increase in SLA for slope trees, whereas populations developed on drier sites at the European scale showed no change in stem hydraulic traits, an increase in *Hv* and a decrease in SLA (Martinez-vilalta et al., 2009; Rosas et al. 2019). This confirms the interest in investigating the tree response over long periods of time and under more constraining conditions, in particular drier and hotter than those encountered in the current range of the species.

### Responses of Scots pines to the harsh conditions of slag slopes

All the slope trees are homogeneous in height and age, indicating that they were installed at the same time, probably when conditions allowed seedlings to settle. This suggests that conditions are mostly too restrictive for Scots pine seedlings to settle and that only the most resistant individuals were able to establish and grow. Therefore, the observed response of the studied trees could be explained by both a selection process of the most resistant genotype and having a phenotypic plasticity adapted to this environment.

The weak growth in slope trees, and to a lesser extent in upslope trees, took place over several years (figure 2). The resulting dwarf phenotype observed on slope trees (figure 1 and 2), and to a lesser extent on upslope trees, would be related to a decrease in parameters related to photosynthetic capacities, such as *K*_s_ and LSC, photosynthetic pigment contents, N content and needle dimensions (table 2 and 3). Since the three conditions are a few meters apart (a few dozen at most), the slope factor would mainly affect water availability and nutrient content (Frain, 1991). According to the results on the nutrient assays (table 1), Scots pines did not suffer from any deficiency for the nutrient assayed, except for N that was at lower content in both slope and upslope trees. This lower N content was already reported as correlated with a decrease in chlorophyll and carotenoids contents and an increase in the chlorophyll a/b ratio (Mu and Chen, 2021). These changes in pigment contents and ratio have also been reported for Scots pines under prolonged drought conditions (Zlobin et al., 2019). This was attributed to a limited capacity to absorb nitrogen (Salazar-Tortosa et al., 2018, Zlobin et al., 2019). Under prolonged drought conditions, Scots pines would be trapped in a feedback cycle of the nutrient deficit that would exacerbate the detrimental impacts of drought on growth (Salazar-Tortosa et al., 2018). It is thus difficult to dissociate nutritional effect from drought effects, especially on long term responses. The vulnerability to embolism is known to decrease in drier conditions (Awad et al., 2010, Lemaire et al., 2021), but the effect of nitrogen availability on the hydraulic traits are contradictory among studies (e.g. Hacke et al., 2010; Plavcová et al., 2012; Borghetti et al., 2017). It is thus difficult to analyze the cause of the phenotypic responses in our study. However, some parameters such as growth and photosynthetic pigment contents indicated that slope trees were more stressed than upslope trees, while they showed the same nitrogen content. This suggests that the phenotype of slope trees would be more associated with a water deficiency.

### Prioritizing hydraulic safety in slope trees

Slope trees, and to a lesser extent upslope trees, showed lower vulnerability to embolism than the downslope trees. Several studies have shown that vulnerability to embolism decreases when plants grow under drier conditions; this trait being designed according to the water potential experienced by the plant at the time of xylem formation (Awad et al., 2010, Lemaire et al., 2021). Yet, Scots pine populations growing in very dry conditions do not show changes in their vulnerability to embolism, but rather acclimation of other traits such as *Hv, K*_s_ and *Ψ*_TLP_ (Martinez-Vilalta, 2009; Rosas et al., 2019). Studies on other species concluded the same lack of relationship between climate dryness and vulnerability to embolism (e.g. Herbette et al., 2010; Wortemman et al., 2011; Hajek et al., 2016). A reduction in the vulnerability to embolism would only be possible for the most marginal populations (Stojnic et al., 2017). This is consistent with the fact that slope pines were in conditions at the limit of their ecological niche. It remains to determine if this results from acclimation or from selection of individuals having the lowest vulnerability to embolism.

Slope pines had also a higher *Ψ*_TLP_, limiting water loss through early stomatal closure. This appears contradictory to the results from Rosas et al (2019) showing a lower *Ψ*_TLP_ for Scots pine populations growing on very dry sites. As for vulnerability to embolism, such discrepancy could be related to the fact that the slope pines were in very stressful conditions, and they should ensure their survival rather than an efficient growth. The resulting safety margin is significantly increased in these slope pines. The simulated dynamics of xylem embolism support this improved safety margin for slope trees since increase in xylem embolism was delayed for slope trees compared to other trees when they were submitted to similar drought conditions (Figure 6B and C). Moreover, the slope trees maintained the lowest *g*_min_ value, whereas *g*_min_ was the highest in upslope trees (table 3). This hydraulic safety showed its effectiveness by very low levels of embolism accumulated over the previous years (figure 5). Moreover, the most surprising point is that these trees do not seem to be in a more constrained water status, as revealed by the 2 years of monitoring of water potentials (figure 3). A recent study on grapevine based on model-assisted ideotyping demonstrated that, among the various trait combinations tested, the most drought plants had generally a larger hydraulic safety margin (Dayer et al., 2022). In our study, the harsh conditions imposed by the slag slopes favored the development of a large hydraulic safety margin.

This greater hydraulic safety in slope pines would be ensured at the expense of the growth, and thus contributes to the dwarf phenotype of the slope pines. First, there is a carbon cost associated with the development of embolism-resistant xylem (Hacke et al. 2001). In a previous study, we demonstrated that a decrease in vulnerability to embolism was prioritized over the growth for beech recovering from a severe drought (Herbette et al., 2021). Accordingly, we observed an increase in wood density for slope pines (data not shown), a trait that is usually correlated with vulnerability to embolism and requires an energy cost (Hacke et al., 2001). The earlier closure of stomata led to a reduction in carbon assimilation which would further explain the poor growth of slope pines, in addition to reductions in chlorophyll content and needle size. The needles of these slope pines also showed an increase in π_0_ and an increase in ε_0_ which revealed decreases in soluble sugar content and wall thickness, related to a lower photosynthetic activity in these pines. The lever to compensate for this drastic decrease in photosynthetic uptake is an increase in SLA for slope pines.

### Conclusions

Our study showed that Scots pine trees grown for several years under harsh conditions, particularly in terms of water availability, showed physiological adjustments that increased their hydraulic safety at the expense of their growth, which was strongly affected. Contrary to our initial hypotheses, the trees did not operate with an higher hydraulic risk and a sub-lethal embolism threshold, but with a lower hydraulic risk than the upslope and downslope pines having a higher growth rate. On the long term, hydraulic safety is prioritized over efficient photosynthesis and growth that just barely allowed for renewal of structures.

This study helps to improve our forecasts on the response of Scots pine populations to climate change that will be accompanied by intense and recurrent droughts over several years. Of course, such a study does not integrate the interactions between species in the response to climate change, and in particular the competition by more xerophilic species. The next step of this work will be to distinguish the role of plasticity from that of a selection effect on these physiological adjustments.

## Supporting information

Exhaustive list of variables used for SurEau simulations

## Supplementary data

Table S1. Exhaustive list of variables used for *SurEau* simulations.

## Conflict of Interest

The authors declare that the research was conducted in the absence of any commercial or financial relationships that could be construed as a potential conflict of interest.

## Acknowledgements

We are grateful to the AgroClim unit for providing meteorological data, to Pierre Conchon and Marc Vandame for field and lab assistance.

## Author’s contributions

HC, SH and STB contributed to developing the question and experimental design. DSFF, DMSP, HC, JN, PH, SH and STB performed ecophysiological analyses. ME and SH handled experiments dealing with nutrient, chlorophyll and carotenoid contents. EB and PC conducted microCT experiments. STB, HC and SH wrote the manuscript with contributions from all authors.

## Notes

### Competing Interest Statement

The authors have declared no competing interest.

